## Supplementary Figures and Information for "Large transient assemblies of Apaf1 constitute the apoptosome in cells"

**This Supplementary material file includes:**
Supplementary Figures S1 to S5
Captions for Movies S1 to S9
References

### Supplementary Figures:

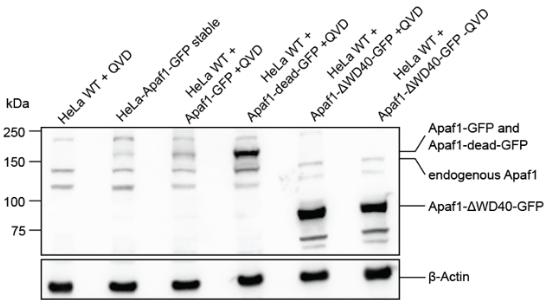


**Suppl. Fig. S1: Apaf1 expression.** Western blot of untransfected HeLa cells, HeLa cells stably or transiently expressing Apaf1-GFP (169 kDa), or transiently expressing Apaf1-dead-GFP (169 kDa) or Apaf1-ΔWD40-GFP (91 kDa) in presence or absence of QVD as indicated, detected using an anti-Apaf1 antibody. Endogenous Apaf1 is expected at 142 kDa. An antibody against β-actin was used for the loading control.

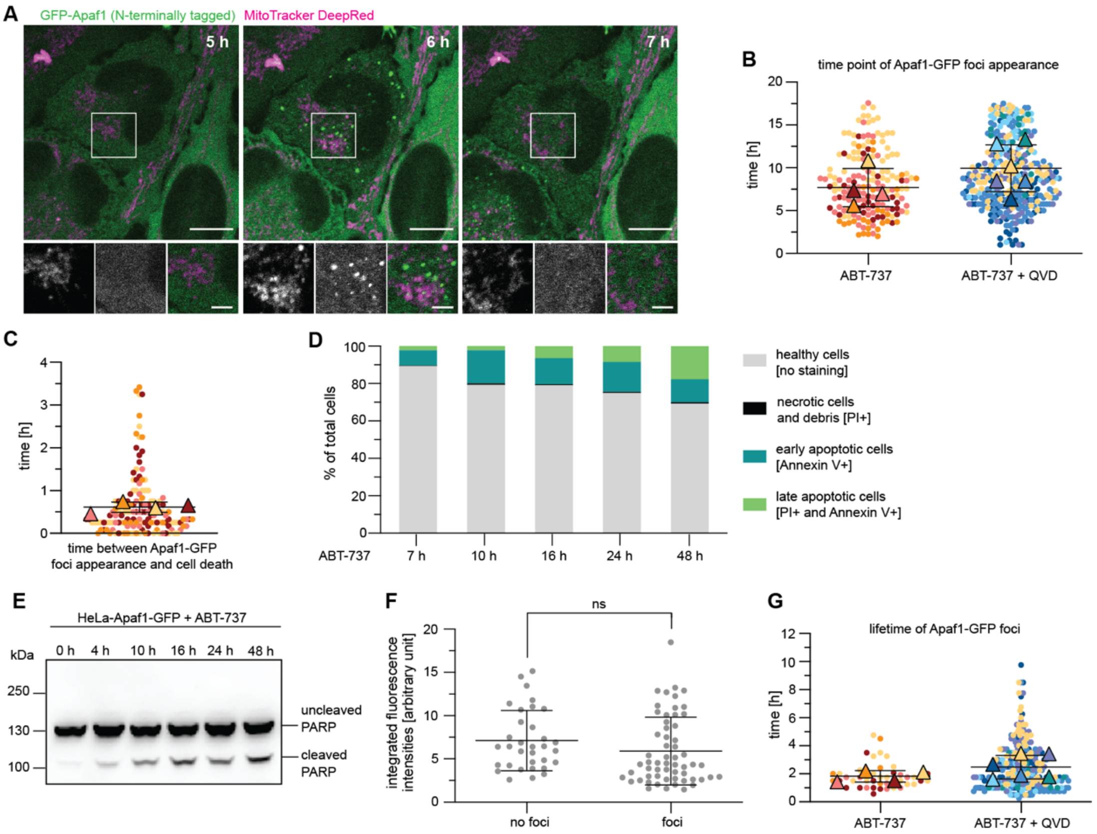


**Suppl. Fig. S2: Characteristics of Apaf1 foci and cell death in HeLa cells. A)** Live fluorescence imaging of HeLa cells transiently expressing a GFP-Apaf1 construct with the GFP tag at the N-terminus, showing Apaf1 foci (green) upon ABT-737 treatment, similarly to the C-terminal constructs that were used throughout the study. Mitochondria were stained with MitoTracker DeepRed (magenta). Image acquisition time since ABT-737 treatment is indicated on large images. White squares indicate areas shown as close-ups (from left to right: MitoTracker DeepRed; GFP-Apaf1; merge). **B)** Time points of Apaf1-GFP foci appearance in HeLa cells stably expressing Apaf1-GFP, since treatment with ABT-737, in presence or absence of QVD. Each dot represents an individual cell. Each colour represents an experiment. Means of each experiment are indicated by triangles. Black lines indicate overall means and SD. With ABT-737: Mean=7 h 42 min, SD=2 h 14 min, N=4 experiments. With ABT-737 and QVD: Mean=9 h 57 min, SD=2 h 44 min, N=6 experiments. At least 17 cells were imaged per condition and experiment. **C)** Time elapsed between Apaf1-GFP foci appearance and cell shrinkage (indicating cell death) in HeLa cells stably expressing Apaf1-GFP, treated with ABT-737. Dots represent individual cells, colour-coded according to experiment. Means of each experiment are indicated by triangles. Black lines indicate overall mean and SD. Mean=37 min, SD=7 min, N=4 experiments. At least 49 cells were imaged per condition and experiment. **D)** Percentages of healthy (grey), necrotic (black), early apoptotic (dark green), or late apoptotic (light green) HeLa cells stably expressing Apaf1-GFP at different time points after ABT-737 treatment, determined by FACS using propidium iodide (PI) and Annexin V staining. N=1 experiment. **E)** Western blot showing poly(ADP-ribose) polymerase-1 (PARP) cleavage, indicative of effector caspase activity, in HeLa cells stably expressing Apaf1-GFP at different time points after ABT-737 treatment, detected using an anti-PARP antibody. **F)** Integrated fluorescence intensities of HeLa cells stably expressing Apaf1-GFP, determined in a frame 10 minutes before shrinkage, indicating cell death (for cells without foci), or 10 minutes before formation of Apaf1 foci (for cells with foci). Dots represent individual cells. Black lines indicate mean and SD, in arbitrary units. Cells without foci: Mean=7117, SD=3484. N=33 cells. Cells with foci:

Mean=5893, SD=3917, N= 58 cells. Same imaging data used as in Figures 1A-E. **G)** Lifetimes of Apaf1-GFP foci in HeLa cells stably expressing Apaf1-GFP treated with ABT-737, in presence or absence of QVD. Each dot represents an individual cell. Each colour represents an experiment (1). Means of each experiment are indicated by triangles. Black lines indicate overall mean and SD. For ABT-737 treated cells: Mean=1 h 49 min, SD=24 min, N=4 experiments. For ABT-737 and QVD treated cells: Mean=2 h 29 min, SD=50 min, N=6 experiments. At least 17 cells were imaged per condition and experiment. Scale bars in B: 10 µm in large images, 3 µm in close-ups.

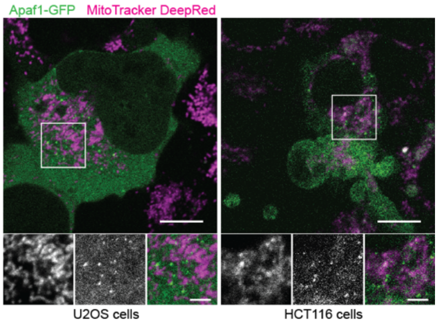


**Suppl. Fig. S3: Apaf1 foci form also in U2OS and HCT116 cells.** Live fluorescence imaging of U2OS and HCT116 cells, transiently expressing Apaf1-GFP (green), showing Apaf1 foci formation upon ABT-737 treatment. Mitochondria were stained with MitoTracker DeepRed (magenta).

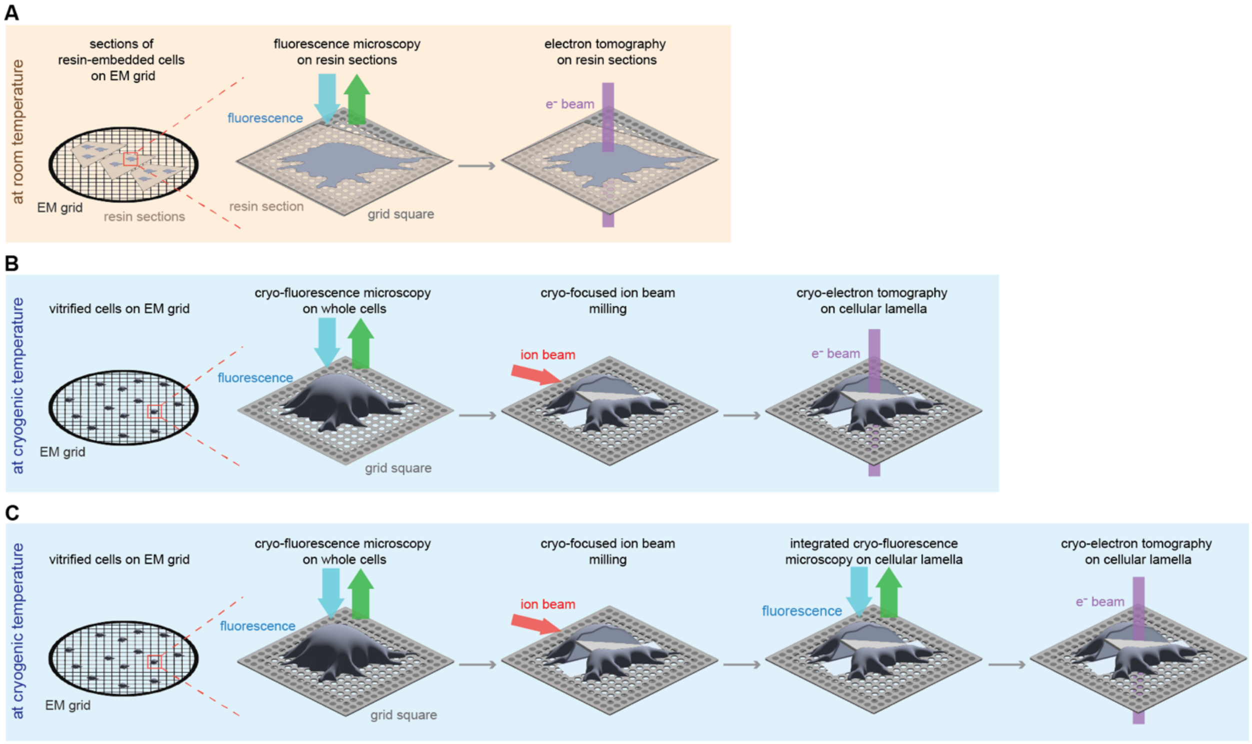


**Suppl. Fig. S4: Correlative light and electron microscopy (CLEM) workflows to identify and image the structural organization of Apaf1 foci in cells.** See also materials and methods for details. **A)** In-resin CLEM (2,3): After high-pressure freezing, freeze-substitution and embedding, samples are sectioned into approximately 200 nm thin sections, which are deposited on electron microscopy (EM) grids. The grids are imaged by fluorescence microscopy to identify regions of the sections containing cell areas with fluorescent signals of interest, here Apaf1-GFP foci. These areas are found back in the electron microscope, and electron tomograms are acquired at room temperature. The position of the tomograms is correlated with high precision to the fluorescence images to determine the centroid positions of the Apaf1-GFP foci. **B)** Pre-focused ion beam (FIB) milling cryo-CLEM (3,4): Cells are grown on EM grids and vitrified by plunge freezing before the EM grids are imaged by cryo-fluorescence microscopy to identify cells containing fluorescent signals of interest, here Apaf1-GFP foci. The EM grids are then transferred to a cryo-FIB scanning electron microscope (SEM), where cells identified to contain Apaf1-GFP foci are thinned to obtain approximately 200 nm thin lamella. The EM grids containing lamella are then transferred into a cryo-transmission electron microscope (TEM) for cryo-electron tomography (cryo-ET). The regions of cryo-ET acquisition are selected based on correlation of the whole-cell cryo-fluorescence images with lamellae overview images. This approach is less precise than the correlation obtained through the workflows shown in A and C. **C)** Pre- and post-FIB milling cryo-CLEM (5,6): Cells are grown on EM grids and vitrified by plunge freezing before the EM grids are imaged by cryo-fluorescence microscopy to identify cells containing the fluorescent signals of interest, here Apaf1-SNAP647 foci. The EM grids are then transferred to a cryo-FIB SEM, where cells identified to contain signals of Apaf1-SNAP647 foci are thinned to obtain approximately 200 nm thin lamella. To confirm presence and precise location of the signals of interest in the lamellae post-milling, the lamellae are imaged using an integrated cryo-fluorescence microscope within the cryo-FIB-SEM (7). The EM grids are then transferred into a cryo-TEM for cryo-ET. The regions of cryo-ET acquisition are selected based on correlation of post-FIB milling cryo-fluorescence images with lamellae overview cryo-TEM images, allowing a precise localisation of Apaf1-SNAP647 foci signals.

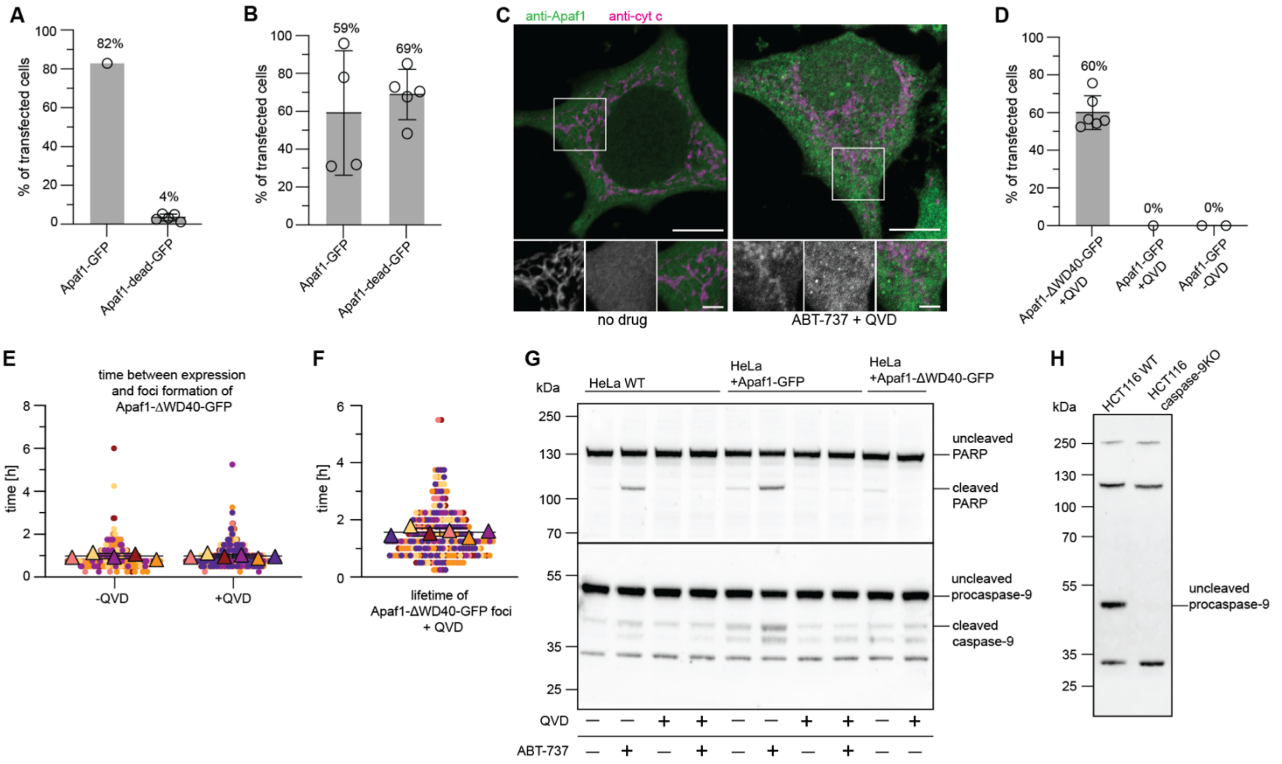


**Suppl. Fig. S5: Apaf1-dead-GFP and Apaf1-ΔWD40-GFP foci formation and characteristics in HeLa cells. A)** Percentages of HeLa cells transiently transfected with Apaf1-GFP or Apaf1-dead-GFP, treated with ABT-737 and QVD, showing Apaf1 foci formation, among all transfected cells. Note that a similar experiment is shown in Figure 4C but without QVD. Lines correspond to mean and SD. Apaf1-dead-GFP: Mean=4%, SD=2%, N=5 experiments. Apaf1-GFP: 82%, N=1. At least 60 cells were imaged per condition and experiment. **B)** Percentages of ABT-737-treated HeLa cells transiently expressing Apaf1-GFP or Apaf1-dead-GFP that shrank, indicating cell death, among all transfected cells. Lines correspond to means and SD. Apaf1-GFP: Mean=59%, SD=33%, N=4 experiments. Apaf1-dead-GFP: Mean=87%, SD=13%, N=5 experiments. At least 68 cells were imaged per condition and experiment. **C)** Immunofluorescence of untreated and ABT-737/QVD-treated HeLa cells, using antibodies labelling cyt c (magenta) and endogenous Apaf1 (green). **D)** Percentages of HeLa cells transiently transfected with Apaf1-ΔWD40-GFP or Apaf1-GFP forming foci without induction of apoptosis by ABT-737, in presence or absence of QVD as indicated, among all transfected cells. Note that a similar experiment is shown in Figure 4F but without QVD. Black lines correspond to mean and SD. Apaf1-ΔWD40-GFP: Mean=60%, SD=9%, N=6 experiments. Apaf1-GFP with QVD: 0% N=1 experiment. Apaf1-GFP without QVD: Mean=0%, SD=0%, N=2 experiments. At least 72 cells were imaged per condition and experiment. **E)** Time elapsed between the first detection of the fluorescent signal of Apaf1-ΔWD40-GFP and foci formation, in HeLa cells in absence or presence of QVD. Dots represent individual cells, colour-coded according to experiment. Mean lifetimes of each experiment are indicated by triangles. Black lines indicate overall means and SD. Without QVD: Mean=1 h, SD=8 min, N=5 experiments. With QVD: Mean=1 h SD=5 min, N=6 experiments. At least 74 cells were imaged per condition and experiment. **F)** Lifetimes of Apaf1-ΔWD40-GFP foci in HeLa cells treated with QVD. Dots represent individual cells, color-coded according to experiment. Mean lifetimes of each experiment are indicated by triangles. Black lines indicate overall mean and SD. Mean=1 h 34 min, SD=9 min, N=6 experiments. At least 74 cells were imaged per condition and experiment. **G)** Western blots showing PARP and caspase-9 cleavage in HeLa WT cells, or HeLa cells transiently expressing Apaf1-GFP or Apaf1-ΔWD40-GFP, in presence or absence of ABT-737 and QVD. The two blots are from one single membrane cut into two; the upper part was incubated with an anti-PARP antibody, the lower part with an anti-caspase-9 antibody. **H)** Western blot showing knockout of caspase-9 in HCT116 cells, detected using an anti-caspase-9 antibody. Scale bars in C: 10 µm in large images, 3 µm in close-ups.

**Movie captions:**

**Movie S1. Live fluorescence imaging of HeLa cells stably expressing Apaf1-GFP, treated with ABT-737.** Movie showing Apaf1-GFP (green) foci formation followed by cell shrinkage and death. Mitochondria were stained with MitoTracker DeepRed (magenta). Image acquisition time after ABT-737 treatment is indicated at the top left. Imaging frame rate: 15 minutes. Scale bar: 10 µm.

**Movie S2. Live fluorescence imaging of a HeLa cell stably expressing Apaf1-GFP, treated with ABT-737.** Movie showing two consecutive events of Apaf1-GFP (green) foci formation in an apoptotic cell. Mitochondria were stained with MitoTracker DeepRed (magenta). Image acquisition time after ABT-737 treatment is indicated at the top left. Imaging frame rate: 15 minutes. Scale bar: 10 µm.

**Movie S3. Live fluorescence imaging of a HeLa cells stably expressing Apaf1-GFP, treated with ABT-737 and QVD.** Movie showing Apaf1-GFP (green) foci formation and disassembly in apoptotic cells. Mitochondria were stained with MitoTracker DeepRed (magenta). Image acquisition time after ABT-737 and QVD treatment is indicated at the top left. Imaging frame rate: 15 minutes. Scale bar: 10 µm.

**Movie S4. Electron tomogram of Apaf1-GFP in resin-embedded cell.** Movie through virtual slices of the electron tomogram shown in Fig. 3B, obtained by CLEM on resin-embedded HeLa cells expressing Apaf1-GFP, treated with ABT-737 and QVD. Scale bar: 200 nm.

**Movie S5. Electron tomogram of Apaf1-GFP in resin-embedded cell.** Movie through virtual slices of the electron tomogram shown in Fig. 3D, obtained by CLEM on resin-embedded HeLa cells expressing Apaf1-GFP, treated with ABT-737 and QVD. Scale bar: 200 nm.

**Movie S6. Cryo-electron tomogram of Apaf1-GFP in vitrified, cryo-FIB-milled cell.** Movie through virtual slices of the cryo-electron tomogram shown in Fig. 3G, obtained by pre-FIB milling cryo-CLEM of HeLa cells expressing Apaf1-GFP, treated with ABT-737 and QVD. Scale bar: 100 nm.

**Movie S7. Cryo-electron tomogram of Apaf1-SNAP647 in vitrified, cryo-FIB-milled cell.** Movie through virtual slices of the cryo-electron tomogram shown in Fig. 3L, obtained by pre- and post-FIB milling cryo-CLEM of HeLa cells expressing Apaf1-SNAP-tag labelled with SNAP-Cell 647-SiR, treated with ABT-737 and QVD. Scale bar: 100 nm.

**Movie S8. Live fluorescence imaging of a HeLa cells stably expressing Apaf1-GFP, microinjected with cyt c.** Movie showing Apaf1-GFP (green) foci formation and dynamics upon microinjection of cyt c and rhodamine dextran (magenta). Imaging time after the start of the microinjection session is indicated at the top left. Imaging frame rate: 15 minutes. Scale bar: 10 µm.

**Supplementary Movie S9. Live fluorescence imaging of a HeLa cells expressing Apaf1-ΔWD40-GFP.** Movie showing Apaf1-ΔWD40-GFP (green) foci formation and disassembly upon expression in cells. Imaging time after the transfection is indicated at the top left. Imaging frame rate: 15 minutes. Scale bar: 10 µm.
